# Candidate SNP analyses integrated with mRNA expression and hormone levels reveal influence on mammographic density and breast cancer risk

**DOI:** 10.1101/259002

**Authors:** M. Biong, M. Suderman, VD. Haakensen, B. Kulle, PR. Berg, I.T. Gram, V. Dumeaux, G. Ursin, Å Helland, M. H Hallett, AL Børresen-Dale, V.N. Kristensen

**Author notes:** Margarethe Biong, Matthew Suderman, Vilde Drageset Haakensen, Åslaug Helland, Bettina Kulle, Paul Berg, Inger Torhild Gram, Vanessa Dumeaux, Giske Ursin, Michael Hallett, Anne-Lise Børresen-Dale, Vessela Kristensen. Address correspondence to: Vessela N. Kristensen, Oslo University Hospital, The Norwegian Radium Hospital, Institute for Cancer Research, Department of Cancer Genetics, Montebello 0310, Oslo, Norway, Tel. ^*^ 47 22 78 13 75.

## Introduction

Mammographic density (MD) is a well-documented risk factor for breast cancer[1–9], and thus a promising target for early detection of the disease. Based on mammographic results, the breast is given a percent density (PDEN) score ranging from 0-100% with 0 being the least dense (primarily adipose tissue). It is estimated that the risk of developing breast cancer is four to six times higher in women with an MD score of at least 75% [2,9] and that one-third of breast cancers occur in patients with PDEN scores of at least 50% [2]. MD is thus a considerably larger risk factor than traditional factors such as early menarche and nulliparity[1].

MD develops as a result of numerous life style factors[10,11] as well as genetic predisposition [12,13] and is greatly influenced by hormonal changes during a woman’s life cycle, spanning from puberty through adulthood to menopause. The change in levels of circulating hormones such as estrogen and progesterone, and their receptors cause variations in the degree of proliferation and differentiation of the breast. Of note, during the postmenopausal years the breast is under the influence of either low levels of hormones derived from adipose tissue or, if hormone therapy (HT) is used, high levels of exogenous hormones. Also, results from studies of monozygotic and dizygotic twins show that MD has a strong genetic component accounting for as much as 30-60% of the variability[12,13]. Mutations and polymorphisms in the genes involved in the development of the breast are of great interest since they could potentially disrupt DNA binding sites and gene splicing or change the amino acid composition of the resulting proteins and influence MD. In fact, certain single nucleotide polymorphisms (SNPs) that affect MD have already been identified [14–17].

In this study, we have selected SNPs in genes belonging to the estrogen signaling pathway or with relevance to MD. A total of 257 SNPs in 165 genes were genotyped via the Sequenom MassARRAY platform and analyzed across a discovery (n=403) and a validation (n=51) dataset of subjects with extensive epidemiological information and serum hormone levels. We also generated mRNA expression profiles from breast tissue with density for a subset of these patients, in order to investigate the downstream effects of the identified genotypes.

## Materials and methods

### Study subjects

In total, the discovery and validation datasets contain 454 healthy, postmenopausal women (Table 1) (Supplementary Table 2 & 3) with associated SNP profiles, MD levels, hormone expression level, and additional clinical-epidemiological data.

**Table 1:** Overview of the study population with a short description and a list of the measured metabolites for each cohort included.

| Data set | n<br>(Total) | n<br>(Anlysed) | Serum analyses | Description |
| --- | --- | --- | --- | --- |
| TMBC | 1041 | 403 | IGF1, IGFBP3,<br>IGFratio, Estradiol,<br>Testosterone, SHBG,<br>DHEA, D4, Prolactin. | Age: 55-71 years, postmenopausal.<br>Negative mammograms, extensive diet<br>questionnaire, menstrual and<br>reproductive factors known. Collected for<br>use in mammographic density study. |
| MDG | 186 | 51 | Estradiol,<br>Testosterone, SHBG,<br>FSH, LH ,<br>Progesterone,<br>Prolactin. | Age: 25-69 years, premenopausal,<br>menopausal and postmenopausal with<br>dense breast and small cancers.<br>Questionnaire on hormone use and<br>reproductive factors. Collected for use in<br>mammographic density study. |
| **Total** | **1227** | **454** |  |  |

<sup>Φ</sup> The Mammography and Breast Cancer study
<sup>φ</sup> Mammographic Density and Genetics

**Table 2:**
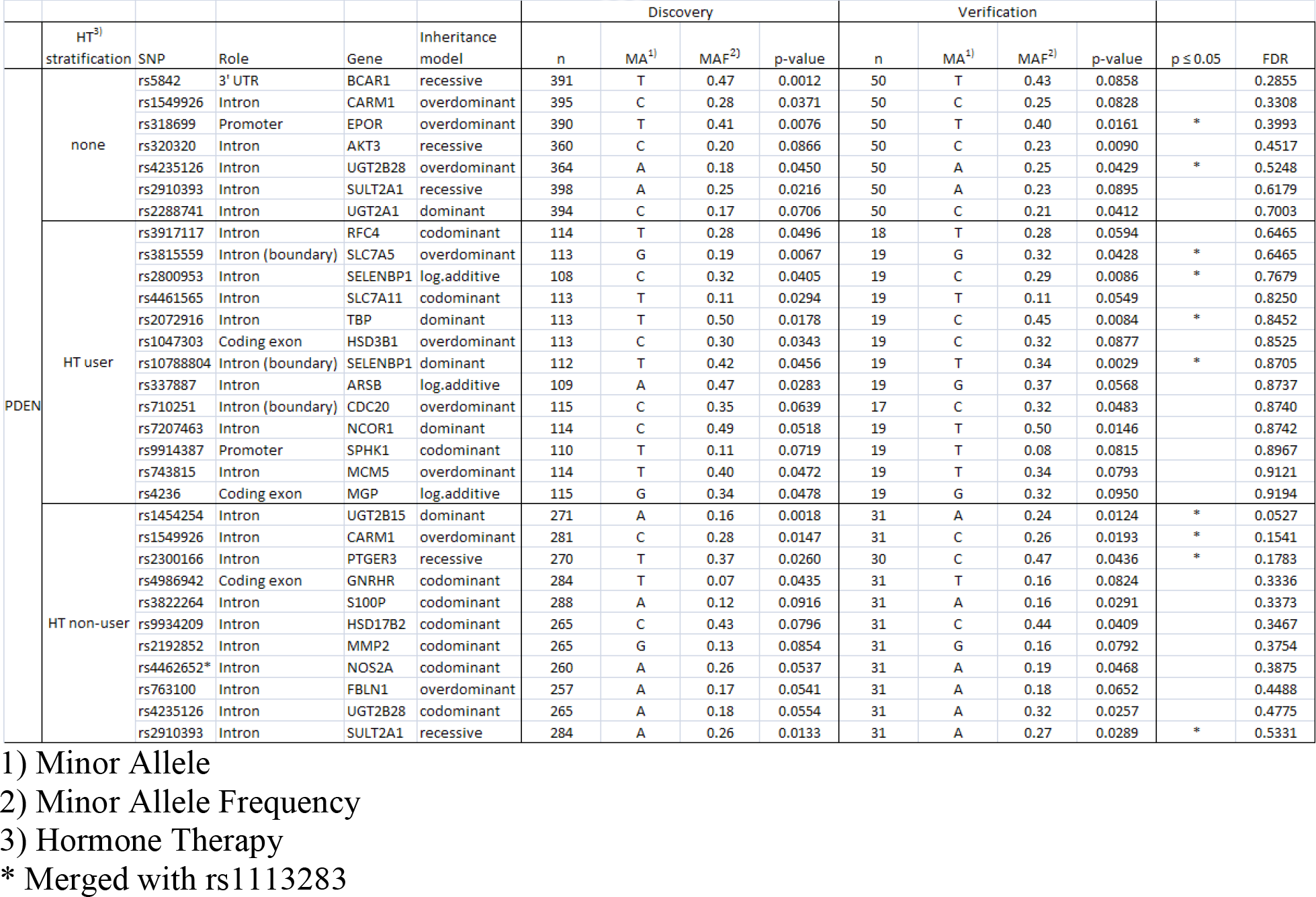
SNPs found significantly associated with PDEN in the discovery and verification set and their overlap according to p-value <0.1, stratified on HT.

**Table 3:** SNPs found associated with testosterone and SHBG in the discovery and verification set and their overlap, HT non-user only.

| Serum<br>hormone | SNP | Gene | Inheritance<br>model | Discovery |  |  |  | Verification |  |  |  |
| --- | --- | --- | --- | --- | --- | --- | --- | --- | --- | --- | --- |
|  |  |  |  | n | MA<sup>1)</sup> | MAF<sup>2)</sup> | p-value | n | MA<sup>1)</sup> | MAF<sup>2)</sup> | p-value |
| SHBG<sup>3)</sup> | rs4986942 | GNRHR | codominant | 279 | T | 0.07 | 0.024 | 32 | T | 0.17 | 0.051 |
| TES<sup>4)</sup> | rs1047303 | HSD3B1 | codominant | 262 | C | 0.26 | 0.054 | 32 | C | 0.33 | 0.040 |

1) Minor Allele
2) Minor Allele Frequency
3) Serum sex hormone binding globulin
4) Serum testosterone

In more detail, (1) the discovery dataset consists of postmenopausal Norwegian women from the Tromsø Mammography and Breast Cancer Study (TMBC) as described elsewhere [18]. In brief, the samples were collected as part of the population-based Norwegian Breast Cancer Screening Program (NBCSP) at the University Hospital of North Norway in the spring of 2001 and 2002. The study consists of postmenopausal women, aged 55-71 years residing in the municipality of Tromsø, Norway. Concomitant with mammography screening, the women were interviewed by a trained research nurse about reproductive and menstrual factors, previous history of cancer, smoking status, and the use of HT or other medications. Measurement of the hormones (estradiol, testosterone, DHAE, vitamin D (D4), prolactin) and glycoprotein (SHBG) were obtained for women not currently using HT. Of a total of 1041 postmenopausal women, 433 were selected for analysis based on the epidemiologic information available. Of these 433, 30 had more than 30% missing genotypes and were removed leaving 403 samples for analysis. All women signed an informed consent and the study was approved by the National Data Inspection Board and the Regional Committee for Medical Research Ethics.

(2)The validation dataset included samples from the Mammographic Density and Genetics (MDG) study as described previously [19]. This dataset contains healthy women (n=120), with low and high MD. The women were recruited either through the Norwegian Breast Cancer Screening Program between 2002 and 2007, or through referral to a breast diagnosis centre for a second look due to irregularities.

Women using anticoagulants, having breast implants or cancer, being pregnant or lactating were excluded. All women underwent mammography and provided information about weight, parity, HT use and family history of breast cancer. Two breast biopsies and three blood samples were collected from each woman. Serum levels of sex hormones were obtained (LH, FSH, prolactin, estradiol, progesterone, SHBG and testosterone). Of these, estrogen, LH and FSH were used to determine menopausal status in combination with age and hormone use (Supplementary Table 4).

Of the 120 samples, we used only the samples for which we obtained genotypes (n=106). Of these, 10 were removed due to lack of clinical information, 45 because they were premenopausal, leaving 51 samples from healthy postmenopausal women for further analysis. All women provided signed informed consent. The study was approved by the local Regional Committee for Medical Research Ethics and local authorities (IRB approval number S-02036).

### Mammographic classifications

The craniocaudal mammogram was digitized for participants in both the discovery and the validation cohorts. Only the left mammogram was obtained for subjects in the discovery cohort, whereas both left and right mammograms were obtained for the validation cohort. A Cobrascan CX-812 scanner at a resolution of 150 pixels per in. was used in the discovery cohort, whereas the Kodak Lumisys 85 scanner (Kodak, Rochester, New York) was used in the validation cohort. MD was assessed by GU and quantified using the University of Southern California Madena computer-based threshold method [20], while the total breast area was assessed by a research assistant trained by GU. Briefly, the method is as follows: the digitized mammogram is viewed on a screen, a reader defines the total breast area using an outlining tool. The region of interest (ROI) is then defined, excluding the pectoralis muscle, prominent veins and fibrous strands. A computer software program is used to determine the pixel value in the image ranging from 0, the darkest shade (black), to 225 the lightest shade (white), with shades of gray being intermediate values. The pixels are then tinted according to a certain threshold representing mammographic densities. The total number of pixels is estimated as well as the number of tinted pixels within the ROI. Absolute density is defined as the count of the tinted pixels within the ROI. Percent density (PDEN) is the absolute density divided by the total breast area multiplied by 100, and is the measurement used for subsequent analyses referred to as mammographic density (MD).

### Gene and SNP selection

Using literature and molecular databases, we selected candidate genes and SNPs within genes that may potentially influence MD. A literature search was completed using Entrez Pubmed [21] with the following keywords: (1)“Estradiol and mammographic density“, (2)“Estradiol and ER”, (3)“SNPs and (1)&(2)”, (4) “Mammographic density”, (5) “Breast density”, and (6)“Single nucleotide polymorphism and Estrogen/Progesterone”. In addition to the selection of genes from the resulting publication lists, we also included genes from the estradiol pathway as defined by CGAP [22] (provided by Biocarta) and iPATH^™^ [23]. PathwayAssist/PathwayStudio®[24] (licensed software by Ariadne Genomics) was used to select additional genes from both the estrogen metabolism and ER signaling pathways. A total of 281 genes were selected through these processes. Candidate SNPs within these with a population frequency greater than 0.01 were subsequently identified using Ensembl [25], SNPper [26] and HapMap [27]. We then analyzed the file from HapMap in Haploview [28] using the De Bakkers “tagger”-test[29] for the identification of correlated SNPs or haplotypes (r^2>0.8). From the sets of correlated SNPs, the software arbitrary identifies one as a representative or tagging SNP, thus removing the need to test the other SNPs in the lists. Lastly we used SIFT [30] (Sorting Intolerant From Tolerant) developed at Fred Hutchinson Cancer Centre in Seattle[31] to reduce our SNP selection to those that might impact protein function based on the sequence homology and physical properties of amino acids. Other SNPs established in the literature to be associated with MD or breast cancer were also included. A total of 1001 SNPs were selected in the 281 genes. Supplementary Figure 1 outlines the gene and SNP selection process.

### DNA isolation and genotyping

The DNA was isolated from blood collected in EDTA-tubes by phenol/chloroform extraction followed by ethanol precipitation using the Applied Biosystems Model 340A Nucleic Acid Extractor and stored in TE-buffer at 2-8°C. Sample concentrations were measured by UV/Vis spectrophotometer (Nanodrop ND-1000) and normalized to 10ng/ul by adding ultrapure water.

All SNPs (n=1001) were annotated and a 200bp flanking sequence at each side of the SNP was extracted from SNPper in the CHIP bioinformatics tools[26] database. The final SNP list was sent to CIGENE at the Norwegian University of Life Sciences (UMB) in Aas for primer design. Genotyping of SNPs was performed using the MassARRAY system from Sequenom (San Diego, USA). The software SpectroDESIGNER v3.0 (Sequenom) was used to design PCR-primers, extension-primers and optimal multiplexes to ensure that little or no unspecific binding occurred due to similar primer design. In total, SNP-assays were successfully designed for 905 SNPs. An initital subset of 519 SNPs was genotype of which 490 SNPs were successful. In quality control we removed SNPs not residing on autosomes (n=9) and that had a call rate of <80% (n=154) or a minor allele frequency (MAF) less than 5% (N=99), leaving 257 SNPs for statistical analysis (Supplementary Figure 2). Multiplex composition and primer sequences are available from the authors on request. All SNP genotyping was performed according to the iPLEX protocol from Sequenom [32]. For allele separation, the Sequenom MassARRAY™ Analyzer (Autoflex mass spectrometer) was used. Genotypes were assigned in real time by the MassARRAY SpectroTYPER RT v3.4 software (Sequenom) based on the mass peaks present. All results were manually inspected, using the MassARRAY TyperAnalyzer v3.3 software (Sequenom).

### Hormone assays

The women in the discovery cohort had non-fasting venous blood samples drawn the day of the mammographic screening. Two 9mL citrate vials were collected for plasma extraction and after centrifugation for 15 minutes at 3000rpm the plasma was stored at -70°C. The hormone analyses were performed at the International Agency for Research on Cancer (IARC). Depending on the hormone, one of the three following methods were used: direct double antibody radioimmunoassays from Diagnostic Systems Laboratories(Webster, TX), direct radioimmunoassay from Immunotech (Marseille, France) or direct “sandwich” immunoradiometric assay from Cis Bio (Gif sur Yvette, France), described elsewhere[18].

In the validation cohort blood was drawn on SST tubes with gel, and then left for 30 minutes on the bench before centrifuging for 10 minutes on 2000 G. Serum was subsequently aliquoted and stored at -20°C. The serum hormone levels were measured with electrochemiluminescence immunoassays (ECLIA) on a Roche Modular E instrument under the supervision of one of the authors (VDH). The laboratory participates in an external quality assessment scheme entitled Labquality, and is accredited according to ISO–ES 17025.

### Gene expression profiling from breast tissue

Biopsies from normal breast tissue were obtained from the validation dataset. Two breast biopsies were taken from each woman with a 14-gauge needle using ultrasound to identify dense areas, in order to avoid purely fatty biopsies. The biopsies were soaked in ethanol (for DNA extraction) and RNA later (for RNA extraction) and were stored at −20°C at Oslo University Hospital. For one hospital the method differed in that the biopsies (92 patients) were snap frozen with liquid nitrogen and stored at −80°C.

The method for the RNA extraction and expression analysis is described in detail elsewhere [19]. Briefly, the homogenization, cell lysis and RNA extraction was performed with the RNeasy Mini protocol (Qiagen, Valencia,CA). Concentrations were determined using a NanoDrop ND-1000 spectrophotometer (Thermo scientific, Wilmington, DE). Amplification and labeling of the RNA was done using the Agilent Low RNA input Fluorescent Linear Amplification Kit Protocol. We used Agilent Human Whole Genome Oligo Microarrays, G4110A, processed by an Agilent scanner via Feature Extraction 9.1.3.1 software (all from Agilent Technologies, Santa Clara, CA). (See supplementary document 13 for gene expression pre-processing and normalization procedure)

### Statistical analysis

The genotype distribution for the study population was assessed for deviation from Hardy Weinberg (HW) equilibrium using the chi-squared (*x*^2^) test (p≤ 0.05). These SNPs were flagged in further analysis (Supplementary Table 12).

#### Genotype and phenotype association

All analyses were performed using R version 2.10.1, apart from the globaltest [33] which was performed in R 2.12.0. To investigate the association between the genotypes and various phenotypes (ie MD, hormone levels and gene expression), we used generalized linear models (GLMs) [34] under five different inheritance models: additive, dominant, codominant, overdominant and recessive. Supplementary Table 3 describes the values of the genotype variable under these inheritance models.

A one-way ANOVA was performed on the discovery dataset and adjusted for age and BMI to estimate the association between MD and each SNP without any reference to inheritance models(results not shown). The p-values from these F-tests, one per SNP, were corrected for multiple testing by calculating the false discovery rate (FDR).

#### Expression and hormone analysis

To discover factors that potentially mediate the effects of SNPs on MD in healthy women, we analyzed microarray expression data from normal breast tissue in the validation dataset as well as serum hormone levels from both cohorts. To be considered a potential mediator, we required that mRNA expression of a transcript was correlated with MD or a given hormone level (partial Pearson’s correlation p≤ 0.05) *in cis* and associated (p≤ 0.05) with at least SNPs. Of the 79 healthy women in the validation dataset, 31 were postmenopausal and had genotyping and expression data available enabling their use in SNP/expression analysis with SNPassoc. Similarly, 35 of the 79 women were postmenopausal and had MD measurements available for MD/expression analysis with partial Pearson’s correlation (Table 4)

**Table 4:**
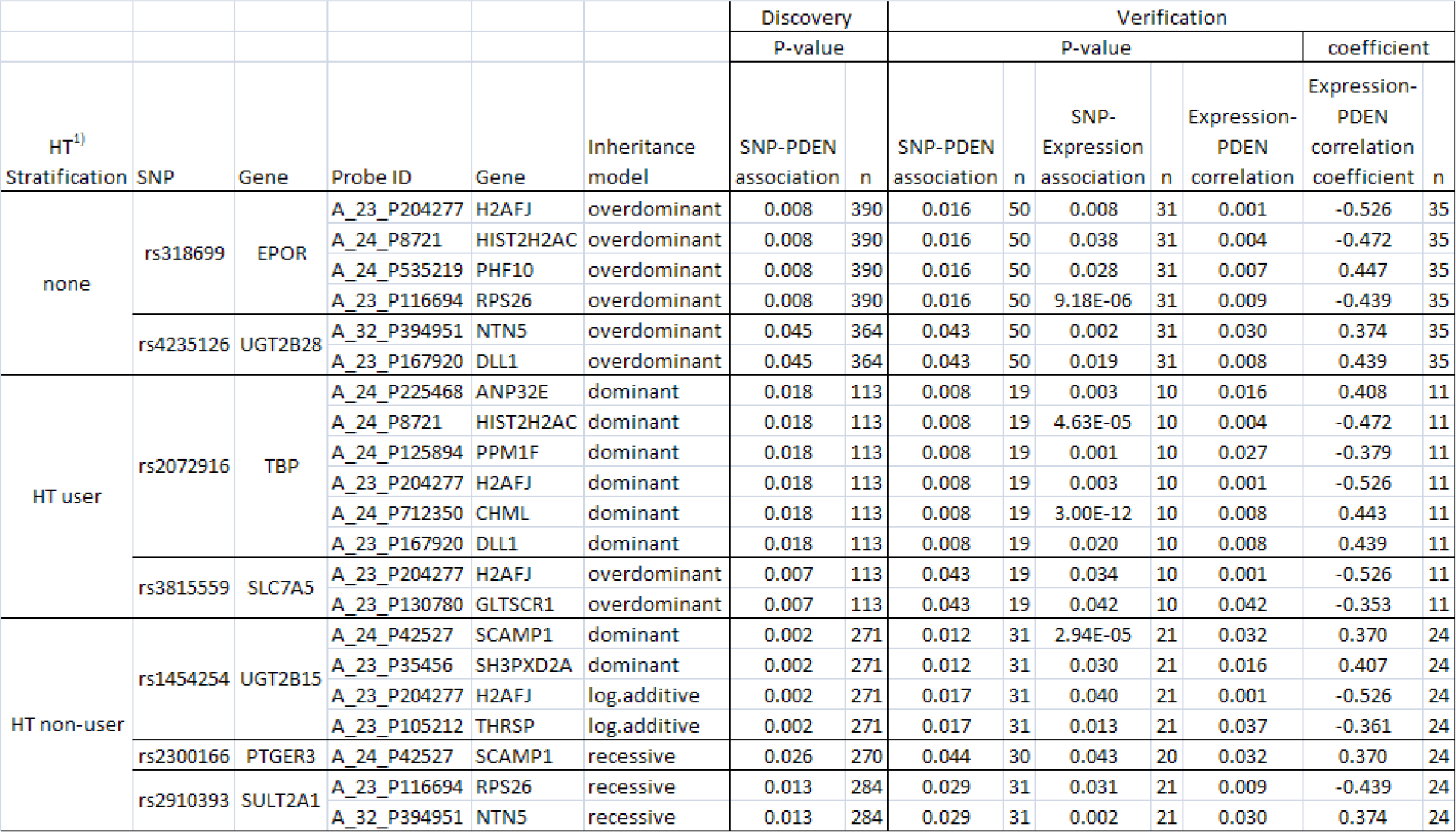
Table listing the significant SNP associations with PDEN (p ≤ 0.05), the correlation of expression with PDEN, and the corresponding correlation coefficient. Results are stratified by HT.

#### Adjustments and stratification

MD decreases with age[35] and is similarly inversely correlated with BMI[36]. High BMI influences MD by way of increased areas of low density due to adipose tissue. We therefore adjusted for these effects in our model. We executed two parallel analyses, one stratifying for HT usage and the other ignoring HT usage. HT users were defined as women who were currently taking HT. We performed separate but identical analyses on all HT strata. We validated the SNPs identified in the discovery dataset associated with MD (p≤ 0.1).

#### Correlations of other variables

Partial correlation of all available variables for each cohort with adjustment for age and BMI was performed independently (Supplementary Tables 5-10).

#### Analysis of SNP sets

The global test [33] was used to determine associations between sets of SNPs and MD. The SNP sets were defined by using results from SNPassoc discovery analysis. For each HT strata two SNP sets were defined: one containing SNPs associated with MD at p≤ 0.05 and the other at p≤ 0.01.

## Results

### Gene & SNP selection

Literature search and database mining yielded 281 genes in the estradiol metabolism, ER and PR signalling pathways. The majority of genes were found by literature search (n=111) and by mining the PathwayStudio^®^ [24]database (n=133). Other tools including CGAP and iPATH provided additional genes (n=12 and 25 respectively). Of the 281 genes we defined 725 haplotype tagging SNPs ( htSNPs) using Hapmap [27] and Haploview [28]. We augmented this set with additional SNPs identified in the literature, and with SNPs located within the 281 genes that are likely to impact protein function, and, with the addition of a few fill-in SNPs (Supplementary Figure 1). The final list consisted of 1001 SNPs residing in 281 genes. Of these, an initial set of 519 SNPs (226 genes) were genotyped on the Sequenom platform. After quality control 257 SNPs (from 165 genes) remained for statistical analyses.

### SNP associations

#### SNPs significantly associated with MD and their downstream effects on serum hormone levels and gene expression

The discovery analyses with and without stratification by HT usage yielded in total 120 SNPs associated with MD at p≤ 0.1 and 77 SNPs at the standard significance level p≤ 0.05 under different inheritance models (results not shown). Of these 120, 31 associations were verified at p≤ 0.1 (no stratification n= 7; HT use n=13; non-HT use n=11, Table 2). Three SNPs overlapped between HT strata, thus 28 unique SNPs were associated with MD across all HT strata. Ten of the 28 SNPs were significantly associated with MD at p≤ 0.05 in both the discovery and validation datasets and are found near genes *EPOR, UGT2B28, TBP, SELENBP1* (2SNPs), *SLC7A5, UGT2B15, CARM1, PTGER3* and *SULT2A1* (Table 2).

Multiple testing correction revealed a relatively high FDR value for the 28 SNPs validated, except within the non-HT stratification. Here three SNPs associated with MD achieved an FDR less than 0.2. SNP rs1454254 in gene *UGTB15* had the lowest FDR in both datasets (FDR=0.053) (Table 2).

Among those 28 SNPs we found two associations with hormones levels in non-HT users: one SNP (rs4986942) in *GNRHR* associated with levels of SHBG (p=0.024 and p=0.051, in the discovery and validation datasets, respectively (Table 3)). The second SNP (rs1047303) in *HDS3B1* was found associated with MD in HT users and with testosterone levels in non-HT users (p=0.054 and p=0.040, in the discovery and validation datasets, respectively (Table 3)). With respect to downstream effects of these SNPs at the gene expression level, seven of the ten SNPs associated with MD at p≤ 0.05 had significant associations with 13 transcripts *in cis* (Table 4). All seven SNPs, of which five were htSNPs, were contained in a unique gene (*EPOR, UGT2B28, TBP, SLC7A5, UGT2B15, PTGER3* and *SULT2A1*).

#### Sets of SNPs associated with MD

In addition to the univariate validation approach described above, we applied a multivariate approach using Globaltest [31]. The SNPsets were defined by the results from the discovery univariate analysis in terms of two p-value thresholds for association with MD (p=0.01 and p=0.05) and stratification by HT use (Supplementary Table 11). For unstratified analyses, seven SNPs near seven genes (SNPset 1) associated with MD (threshold p≤0.01) were validated with a Globaltest p-value of 0.0379 (Table 5). These seven included three SNPs from the univariate analysis. In women currently using HT, 43 SNPs near 36 genes (SNPset 2) associated with MD (threshold p≤ 0.05) were validated with a Globaltest p-value of 0.034 (Table 5). Ten of these 36 were also validated univariately. For HT non-users none of the SNPsets were validated (Table 5). Figure 1 depicts the localization of the genes harbouring the SNPs found associated with MD according to the estradiol pathway.

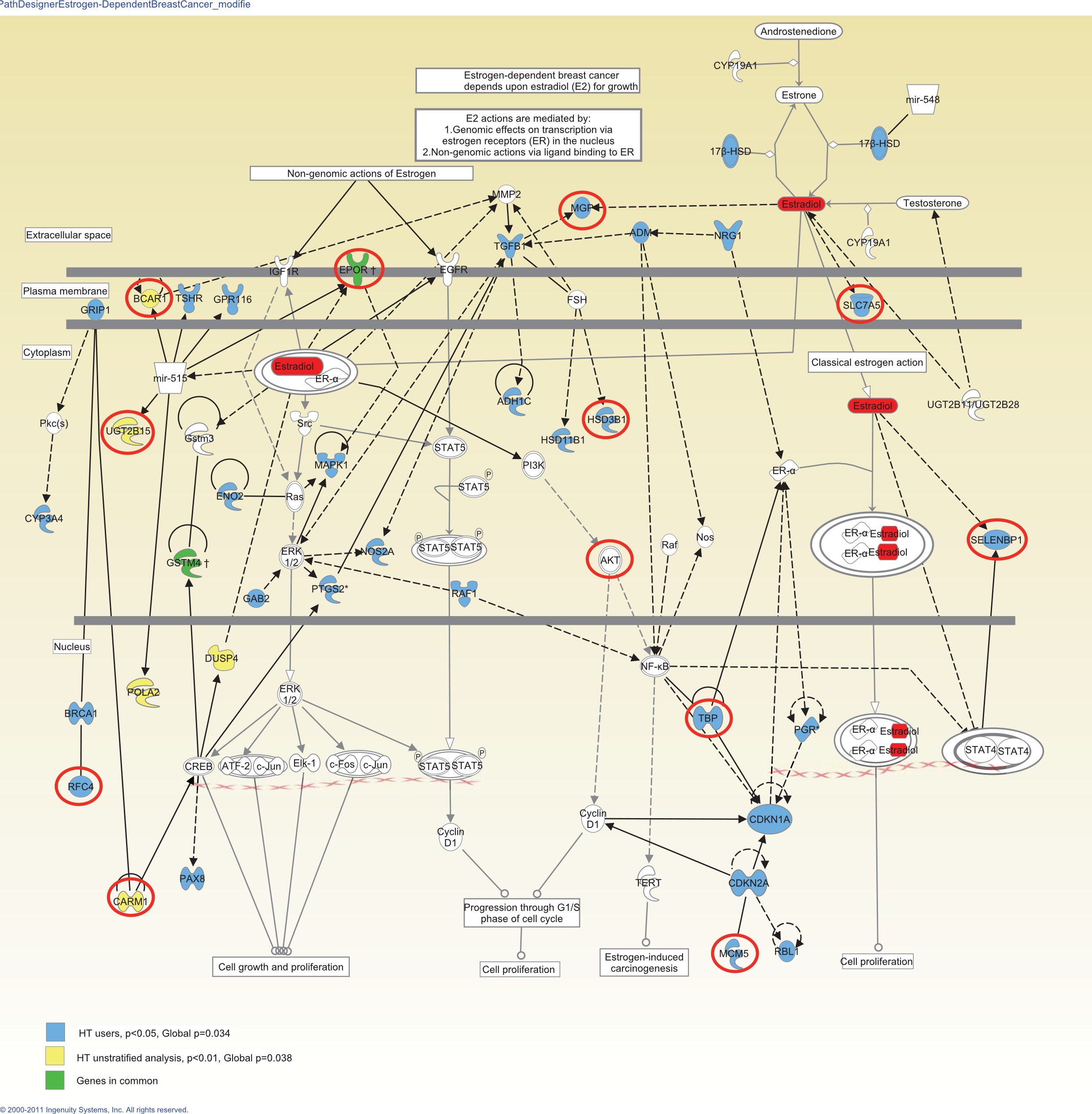

**Table 5:** Globaltest results based on SNPsets defined from results in SNPassoc discovery analysis and validated in the verification dataset.

| HT strata | SNPset | Global p-value | samples(n) | Global p-value | samples(n) |
| --- | --- | --- | --- | --- | --- |
|  | p-value threshold | discovery |  | Verification |  |
| none | $\leq 0.01$ | 3.07E-06 | 403 | 0.038 | 50 |
| | $\leq 0.05$ | 3.01E-11 | 403 | 0.243 | 50 |
| HT users | $\leq 0.01$ | 5.73E-07 | 115 | 0.376 | 19 |
| | $\leq 0.05$ | 5.76E-17 | 115 | 0.034 | 19 |
| HT non users | $\leq 0.01$ | 2.54E-05 | 288 | 0.4 | 31 |
| | $\leq 0.05$ | 1.75E-13 | 288 | 0.43 | 31 |

## Discussion

We have utilized a candidate gene approach in an attempt to clarify the involvement of the estradiol pathway, through the use of SNPs, in MD. Claims about associations of estradiol with MD have been conflicting with some reporting no associations [39–41] and others reporting slight associations [42]. Here we provide evidence for an association with MD by identifying SNPs associated with MD likely to affect the activity of estradiol pathway genes.

Beyond the estradiol pathway, we also obtained results providing evidence for a causal link between MD and BC via SNPs. We showed that some of these associated SNPs were also associated with circulating levels of estradiol, testosterone and SHBG. Finally, although most of the candidate genes were selected from the estradiol pathway, we found SNPs associated with MD that resides in genes that are mainly involved other cancer-related pathways such as signalling, cell cycle including the sex hormone biosynthesis pathway.

### Genes involved in signalling may affect MD

EPOR plays a major role in signalling and upon ligand binding activates the JAK/STAT and Ras/map PI3K/Akt signalling cascades [43–45]. Several studies have implicated EPOR in breast cancer[44,46]. Shi and colleagues showed that EPO, the ligand of EPOR, is able to stimulate phenotypic changes in breast cancer cell lines, and possibly induce metastasis through cell signaling[44]. The SNP (rs318699) in *EPOR* have previously been associated with polycythemia vera (PV), a disorder where blood marrow produces excessive blood cells[47]. Our results suggest that it may be linked to the development of mammographic density.

The *EPOR* SNP was found associated with the expression of *H2AFJ, HIST2H2AC* and *PHF10.* The histone family members *H2AFJ* and *HIST2H2AC* are important in processes involving DNA transcription, repair, replication and stabilization[43]. Also central in transcription regulation is *PHF10*, a gene necessary for multipotent neural stem cell renewal and proliferation[43]. The heterozygote genotype (T/C) of the *EPOR* SNP is associated with decreased expression of *H2AFJ* and higher levels of MD, thus there is an inverse relationship between MD and the expression of *H2AFJ*. This is in support of our previous findings where we found the expression of this gene to be down-regulated in high MD [19]. *H2AFJ* has been proposed to be an oncogene in breast cancer after it was found amplified and over-expressed in breast tumours [48,49]. Similarly, the expression of *HIST2H2AC* was inversely associated with MD, and the heterozygote (T/C) was associated with low expression. To our knowledge no other study has linked this gene to MD or to BC. The heterozygote genotype of rs318699 was associated with increased expression of *PHF10* and increased MD. Banga et al. 2009 reported that PHF10 function is required for cell proliferation[50] thus we hypothesise here that *PHF10* could possibly contribute to higher MD through increased cell proliferation in the breast.

Downstream of EPOR we also identified a SNP in *AKT3* associated with MD. AKT3 plays a role in cell survival, growth factor initiated signalling (e.g. insulin), angiogenesis and tumour formation[43]. Although the verification of the *AKT3* SNP is inconclusive, we observe a strong trend towards involvement of this pathway in the development of MD.

### Members of cell cycle processes may affect MD

The SNPs in *CDC20* and *MCM5* were associated with MD in postmenopausal women currently taking HT. Such women likely have elevated estrogen levels. Estradiol(E2) has previously been found to up-regulate *MCM5*, suggesting participation in DNA stability during increased cell proliferation by E2 [51]. To our knowledge *CDC20* has not been associated with hormones previously but could share the same qualities as *MCM5*.

### Hormone synthesis and clearance in relation to MD

We found seven SNPs in genes *UGT2A1, UGT2B15, UGT2B28, SULT2A1, HSD3B1* and *HSD17B2*. These genes are all central in biosynthesis or clearance of hormones in addition to other substances of special interest is the *UGT* gene family which is involved in the removal of toxic and endogenous substances such as hormones from the human body through glucuronidation. We have previously shown that women under the influence of estrogen, i.e. premenopausal or HT users, have low expression of *UGT2Bs* and high MD[19]. Our current finding further suggests the existence of SNPs in the UGT gene family that may have influence on the development of mammographic density.

The association of the SNP in UGT2B15 (Table 4) with the expression of *SCAMP1, SH3PXD2A* and *THRSP* is of particular interest since expression of *SCAMP1* and *SH3PXD2A* was found positively associated with MD in women not using HT. *SCAMP1* is a carrier protein proposed to participate in endocytosis [52], and *SH3PXD2A* is required for podosome formation and is found in normal immune cells and activated endothelial cells. Podosomes degrade the extracellular matrix and promote cell invasion and are thus suggested to be important in tumor invasiveness, motility and metastasis[53]. Hypothetically the degradation process of the extracellular matrix starts already in normal breast tissue by manifesting in or giving rise to mammographically dense tissue. Lastly, expression of *THRSP* was associated with the *UGT2B15* SNP and inversely associated with MD in women not using HT. Postmenopausal women not using HT can be presumed to have undergone a normal involution and present with breast tissue mainly consisting of type 1 lobules and adipose. *THRSP* has been shown to be expressed in adipocytes, particularly in lipomatous modules. The gene has also been found expressed in lipogenic breast cancers, which suggests a role in controlling tumor lipid metabolism[43]. The *THRSP* association with breast cancer as well as adipose tissue could be explained by Kinlaw et al. who argue that the fatty acids are not carcinogenic but act to fuel metastases[54]. The association between the expression of *THRSP* and the SNP in *UGT2B15* could be explained through their shared influence by lipids, either through lipid metabolism or sex hormone biosynthesis. It is therefore not surprising that we find the expression of *THRSP* associated with MD in these women suggestive of increased expression in breasts mainly consisting of adipose tissue and low MD.

Similarly to the UGT family, *SULT2A1* encodes a protein involved in the metabolism of drugs and endogenous compounds such as hormones and ensures clearance of these compounds and limits their availability. We find here a SNP in *SULT2A1* associated with MD in postmenopausal women not currently taking HT. To our knowledge no association between SNPs in *SULT2A1* and MD has previously been reported. The implications of SULT2A1 in breast cancer mainly involves the regulation of hormone levels present in tumor tissue[55,56]. Upon analysis of expression data we found this SNP associated with the expression of *RSP26* and *NTN5* of which neither has been previously implicated in MD or BC. While little is known about ribosomal protein *RPS26*, it is known that *NTN5* codes for netrin-1-like protein which is involved in axon guidance. Netrins may also act as growth factors by encouraging cell growth activities in target cells. Specifically, netrin1 has been found to be a vascular mitogen, stimulating proliferation, inducing migration and promoting adhesion of endothelial cells and vascular smooth muscle cells[57], which could be of importance in MD.

Another regulator of hormone biosynthesis is the product of *HSD3B1* which catalyses the conversion from progneolone to progesterone. Due to its importance it has been studied in both breast cancer and mammographic density. The SNP rs1047303 has been found to be associated with MD in two independent studies [58,59]. Our findings confirm the association of the variant type allele (Thr) with lower MD compared with the wild type allele (Asn). In addition, we found an association between this SNP and testosterone levels, a novel finding to our knowledge. A previous study found an association of the SNP with the levels of aldosterone [60]. Similarly we found a SNP in *HSD17B2* associated with MD. This gene catalyzes the conversion from estradiol to estrone thus protecting the breast tissue from estradiol. The plots of the genotype distribution in the two materials show conflicting association with MD (data not shown), but based on previous studies and a good p-value in the discovery dataset, these SNPs could potentially affect MD.

### Other SNPs associated with MD

In HT unstratified analysis we identified additional associations not mentioned previously. Of special interest is the association of a SNP in *BCAR1* (breast cancer antiestrogen resistance1) with MD. *BCAR1* is involved in cellular events such as migration, survival, transformation and invasion[43,61], all of which are important in both MD and BC. In addition, high expression levels of *BCAR1* have been linked with poor relapse-free survival and overall survival, and confers a greater risk of resistance to antiestrogen therapy such as tamoxifen [62]. This gene is located on 16q which has been implicated in many cancers including breast cancer[63,64]. We find that being homozygote for the T allele is associated with increased MD, which is to our knowledge a novel finding.

Among the women not taking hormone therapy we find SNPs with validated association with MD in the genes *MMP2* and *NOS2A* (*UGTB15* and *SULT2A1* previously) Of special interest is *MMP2* (matrix metalloproteinase 2) which belongs to a family involved in programmed degradation of extracellular matrix during reproduction, tissue remodelling, arthritis and metastasis[65]. In particular the enzyme encoded by this gene is involved in angiogenesis, tissue repair, tumour invasion and inflammation[43], cancer cell growth, differentiation, apoptosis, migration and immune surveillance [66]. In addition it degrades type IV collagen, which is the major component in basement membranes that underline the epithelium [65]. Associations of MMPs and cancers are numerous and includes cancers of the lung, head and neck, colorectal and hormone related cancers such as, prostate, breast, ovary and endometrium [67]. Associations of single nucleotide polymorphisms inn MMPs with cancer have also been reported[68], but for breast cancer the results are less conclusive[69–71]. We identify here a SNP (rs2192852) in *MMP2* associated with MD in postmenopausal women not taking HT. This is an interesting gene in terms of mammographic density due to its metaplastic ability and could potentially connect MD with BC, and thus the critical point of transition from healthy to a malignant state.

## Globaltest

With the use of Globaltest we identified and validated the association of two SNPsets with MD. This result indicates that the SNPs may hold more potential in a set of SNPs than as single markers.

### Limitations and strengths

These findings support the involvement of genetic variation in genes belonging to the estradiol pathway in a premalignant process, with origin in dense mammographic tissue. There are however limitations to this study. While the discovery set is, to our knowledge, a pure control dataset with a homogenous group of women, the verification dataset is more heterogeneous, including samples from women who have had a suspicious mammogram and can be therefore considered at increased risk of developing breast cancer.

The stratification of the women into HT non-users, never users and past users may also not be optimal since the past user group may include women who had recently stopped using HT. However, we are confident that this might be true for only a few women and that the majority of the HT non-users are representative for women not under the influence of HT.

Our study considered only a few hundred candidate SNPs thus other SNPs associated with MD are missed. However, the SNPs selected were mainly htSNPs with one in linkage disequilibrium (LD) with as many as 65 SNPs, thus it could be said that we tested LD blocks as opposed to single SNP for association with MD, the causative marker(s) may therefore be in LD to the SNPs found here. Of the 257 SNPs, 89 are on the 660GWAS Illumina array. Of the 28 reported with associations here, 11 are on the 660 Illumina array. None of these 11 SNPs were found significantly associated with MD in a 660GWAS study by Lindstrøm *et al.*[72]. SNPs with lower penetrance need a greater sample size due to the stringent significance level applied when testing 100K - 1M SNPs. A candidate gene approach with association to mediatiors and proxymal phenotypes such as mRNA and protein expression decreases this threshold and increases the chance of detecting aberrations with low risk. We did not apply bonferroni correction in the discovery SNP/MD association analysis in SNPassoc. In our study, however, these SNPs were retained and validated in an additional cohort. Agreement of several of our findings with previous studies suggests that they should be further investigated in larger sample populations.

## Conclusion

We find here associations of SNPs within genes involved in the metabolism and regulation of estrogen, a finding that strengthens the existing hypothesis of hormone dependency in MD. We have provided evidence that some of these SNPs may exert an effect on MD via changes in expression levels of certain genes, Further functional studies are required to determine the exact effects of these or other SNPs in LD.

## Abbreviations

MD: mammographic density
BC: breast cancer
HT: Hormone Therapy
BMI: body mass index
SNP: single nucleotide polymorphism

## Authors contributions

The study was supported by grants to VNK from the Norwegian cancer society and the South Eastern Norway Regional Health Authority. MB was supported by grants from the Norwegian Research Council and the South Eastern Norway Regional Health Authority. IT is responsible for collecting TMBC data. ÅH is responsible for collecting MDG data. MB, PRB performed the genotyping lab work. VDH performed the expression array lab work and data processing, assisted in MDG data collection and estimation of mammographic density. GU is responsible for the mammographic density data for both cohorts used.VHD assisted in measuring the serum hormones in MDG samples. MB and MS and BK performed the statistical analysis. MB, MS, and VNK, VD and MH interpreted the results, and MB and MS wrote the paper. VD critically revised and commented on the manuscript. MH provided statistical advice and critically revised and commented on the manuscript. All authors were involved in reviewing the report.

